# Genetic variations associated with adaptation processes in Acrocomia palms: A comparative study across the Neotropic for future crop improvement

**DOI:** 10.1101/2024.08.15.608149

**Authors:** Jonathan A. Morales-Marroquín, Brenda Gabriela Díaz-Hernández, Suelen Alves Vianna, Alessandro Alves-Pereira, Carlos Eduardo de Araújo Batista, Carlos A. Colombo, José Baldin Pinheiro, Maria Imaculada Zucchi

**Author notes:** **Correspondence: Jonathan A. Morales-Marroquín** **Maria I. Zucchi**.

## Abstract

Population genetic research has evolved, focusing on selection processes using single nucleotide polymorphisms (SNP) genotyping techniques to study crop traits and domestication. This study explores the adaptation process of three neotropical palms of *Acrocomia*, a genus that has high potential for oil extraction. Our research focuses on their genetic structure, evolutionary significance, and implications of the selection signatures for breeding efforts. We employed Genotyping-by-Sequencing (GBS) with a focus on outlier SNP markers for identifying adaptive genetic processes in *A. aculeata*, *A. totai*, and *A. intumescens* across the Neotropic. Our results reveal two major gene pools in *A. aculeata*, a North American and a South American group mainly influenced by dispersal and biogeographic barriers, with putative selective signatures linked to fatty acid and carotenoid biosynthesis, pathogen resistance, and environmental stress adaptation. *A. totai* presented a pronounced genetic structure, with SNPs under selection indicating a recent diversification driven by climatic and geological factors, especially in the Pantanal biome. *A. intumescens* displays genetic structuring shaped by the endemic process of biogeographic barriers within the Caatinga biome, with potential shared ancestry with *A. aculeata.* Correlations between allele frequencies and climatic variables highlight adaptation to diverse environments, principally semi-arid, with the annual mean temperature being one of the most influential. Candidate genes associated with fatty acid and carotenoid biosynthesis, as well as pathogen resistance and drought tolerance, indicate targets for domestication. Despite challenges in reduced representation sequencing, this study underlines the potential of *Acrocomia* as a novel crop, offering prospects in oil production, biofuels, and sustainable agriculture. Future efforts should prioritize whole-genome sequencing and genotype-environment interaction studies to realize the full potential of *Acrocomia* in sustainable development and renewable energy production.

## Introduction

The Neotropical palm genus *Acrocomia* (Arecaceae: Arecoideae: Cocoseae: Bactridinae subtribe) holds significant agricultural potential. This genus comprises nine species with diverse distributions across tropical and subtropical America, and is characterized by its solitary, spiny trunk (stipe), persistent remnants of fallen leaf sheaths, and a yellow, coccoid drupe rich in oil [1]. Within the group, *Acrocomia aculeata* (Jacq.) Lodd. ex Mart. presents the widest distribution and greatest economic importance, also *A. totai* Mart. and *A. intumescens* Drude. show promising potential for oil production and bioeconomy [2]. The species are popularly known as Macauba, Coyol, Corozo, Macaw palm (*A. aculeata*), Bocaiuva (*A. totai*), and, Macaiba (*A. intumescens*) [3]. It is worth mentioning that *A. aculeata* and the African oil palm (*Elaeis guineensis* Jacq.) exhibit analogous oil yields and fatty acid compositions [4].

*Acrocomia aculeata*, *A. totai*, and *A. intumescens* can be distinguishable by their distribution and distinct trunk (stipe) traits (Fig. 1 and 2). Wild populations of these palms have exhibited genetic and phenotypic variations linked to environmental adaptability [5–7]. *A. aculeata* is characterized by its spiky trunk and persistent leaf bases from fallen leaves. Its yellow fruits have a diameter ranging from 3.0 to 5.0 cm [4]. The total yield obtained (dry basis) in the extraction of *A. aculeata* fruit pulp oil was 65% w/w from populations of the Brazilian state of Minas Gerais [4], and 53.6% from populations of Costa Rica [8]. Its distribution ranges from the subtropical to tropical regions of northern Mexico through Central America (Guatemala, El Salvador, Honduras, Nicaragua, Costa Rica, and Panama), Caribbean Islands, into Colombia, Venezuela, Guyana, Brazil, and northern Argentina [1]. It is present in six of the seven proposed Neotropical palm biogeographic areas, predominantly in transitions between macro ecoregions like: tropical rainforests (moist broadleaf forest) to drier ecosystems like savannas, grasslands, dry broadleaf forests, and xeric shrublands (Fig. 2) [9,10]. It occurs in the following local biomes: Brazilian Cerrado, Atlantic coastal forest, Amazon, Central America moist forest, Caribbean dry forests, and Llanos [11,12].

**Fig 1.**
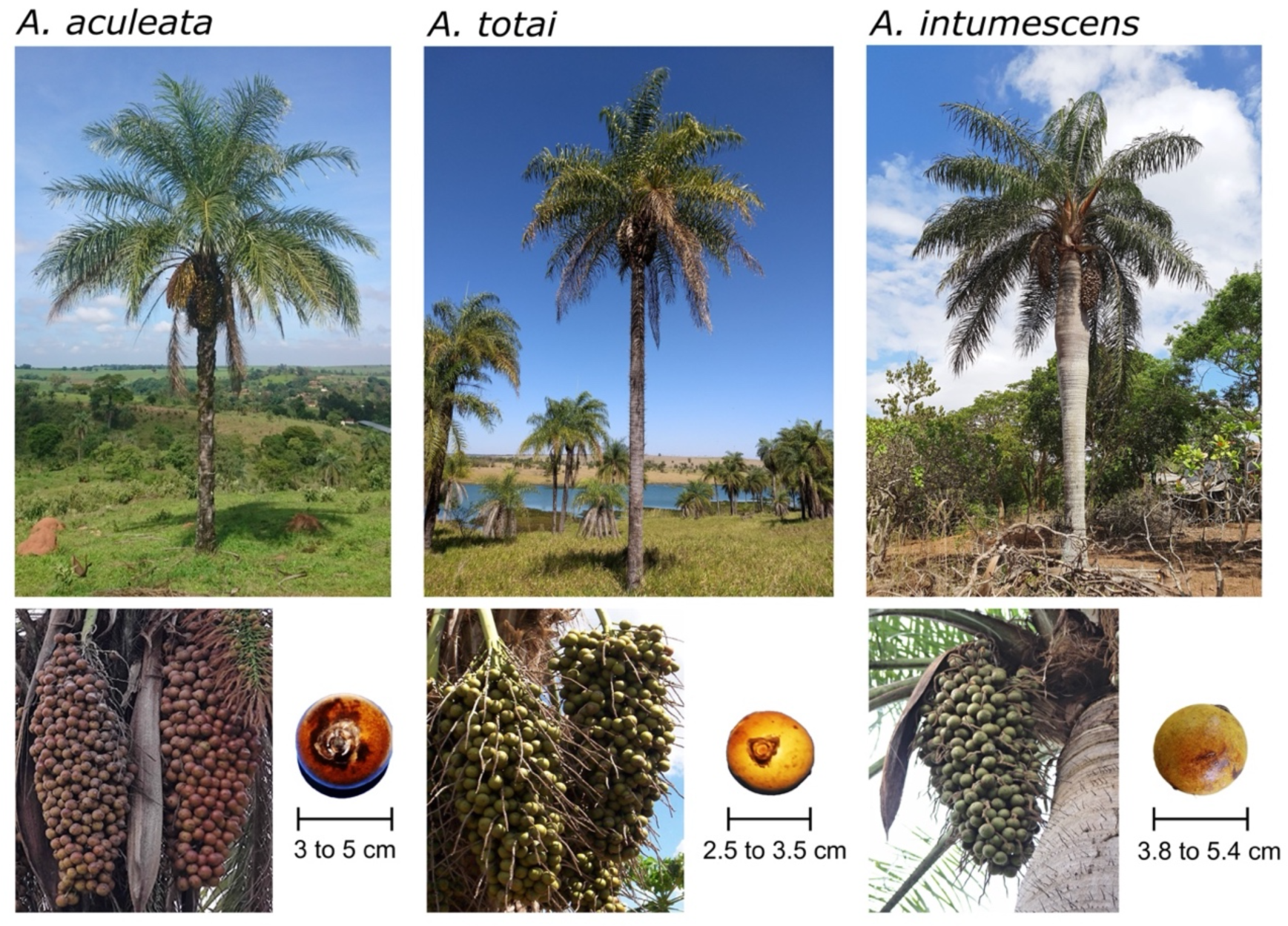
Adult plants of *Acrocomia aculeata*, *Acrocomia totai*, *Acrocomia intumescens*, and their fruits. Location: *A. aculeata* – Palm from Itapira, São Paulo, Brazil. Fruits from Minas Gerais, Brazil. *A. totai* – Palm from Presidente Epitácio, São Paulo, Brazil. Fruits from Corumbá, Mato Grosso do Sul, Brazil. *A. intumescens* – Palm and fruits from Vale da Neblina, Paraíba, Brazil. Photo credits: Brenda Díaz-Hernández (Palms: *A. aculeata* and *A. totai*), Suelen Alves Vianna (Fruits: *A. aculeata* and *A. totai*), and Eulampio Duarte (*A. intumescens* Palm and Fruits).

**Fig 2.**
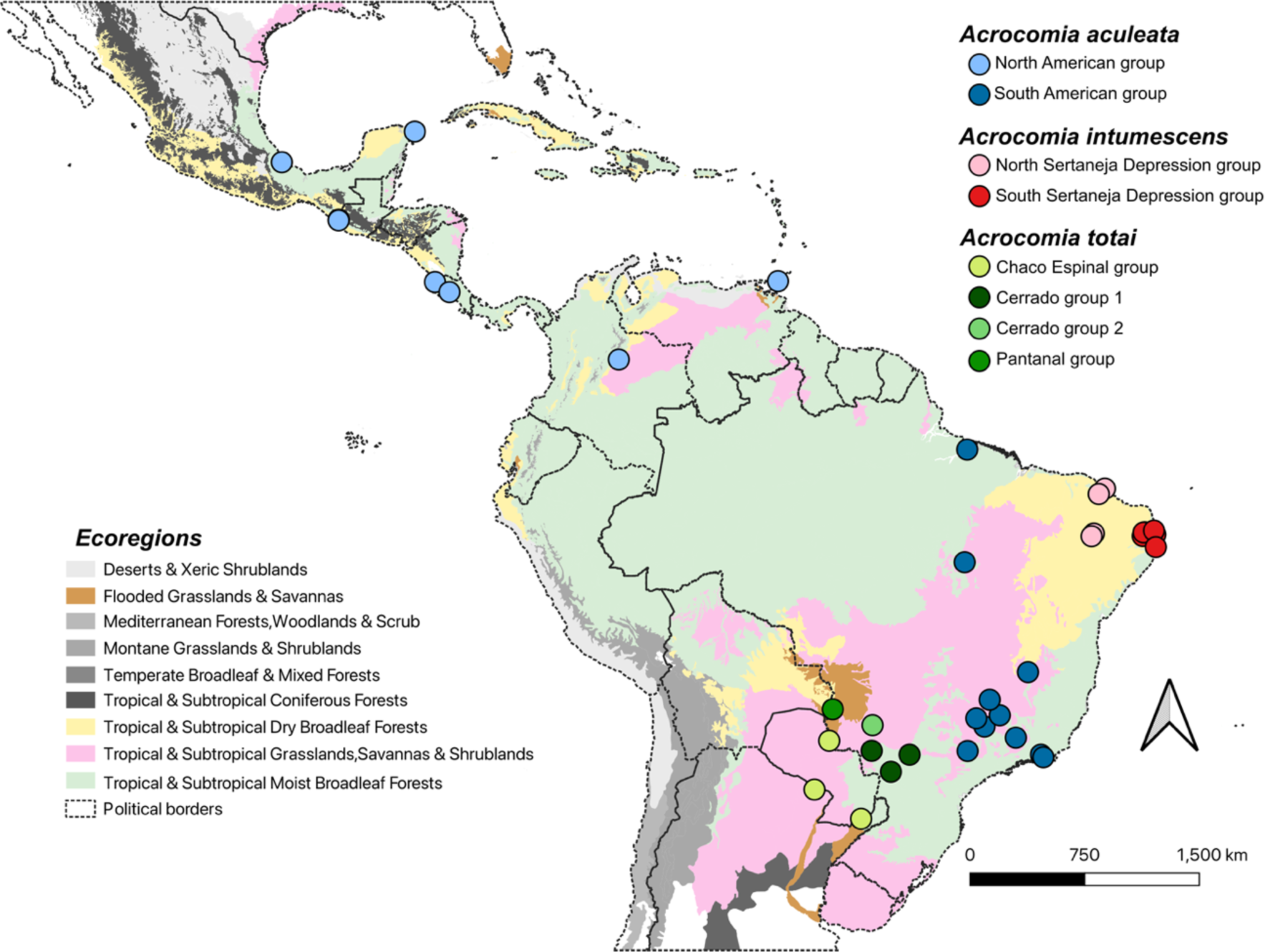
Map showing sampling locations of *Acrocomia* natural populations in the various macro ecoregions of the Neotropic. The map displays populations and major genetic groups within them of *Acrocomia aculeata* (blue dots), *A. totai* (green dots), and *A. intumescens* (red dots), as described in Table 2. The different macro ecoregions of the Neotropic are defined according to Dinerstein et al. [10].

*A. totai* generally features smooth trunk, occasionally with spikes, and rarely covered in persistent leaf bases. The diameter of its yellowish orange fruits ranges from 2.5 to 3.5 cm (Fig. 1) [13]. The total yield obtained (dry basis) in the extraction of *A. totai* pulp oil was 34.70% w/w from populations of the Brazilian state of Mato Grosso do Sul [14]. It is distributed across a preeminent subtropical region in southern Brazil, northern Argentina, Bolivia, and Paraguay, occurring in the local biomes of Brazilian Pantanal, Chaco, and Espinal, as well as the Brazilian Cerrado transition biome. In macro-ecoregion classifications, it occurs in moist broadleaf forests, flooded grasslands, and savannas, as well as in drier ecosystems such as savannas, grasslands, and shrublands (Fig. 2) [10,12]. *A. intumescens* produces yellow fruits with a diameter ranging from 3.8 to 5.4 cm. In its early life, the palm has a swollen trunk with spikes when young, which later becomes smooth in adulthood (Fig. 1) [1]. The total yield obtained (dry basis) in the extraction of *A. intumescens* pulp oil was 26.90% w/w from populations of the Brazilian state of Pernambuco [15]. *A. intumescens* has a more restricted distribution, endemic to the arid and semi-arid tropical regions of the transition zones between the local biomes of Brazilian Caatinga and Cerrado, also in Atlantic coastal forest in the northeastern states of Pernambuco, Ceará, Paraíba, and Maranhão [11]. In macro-ecoregion classifications, it occurs in dry broadleaf forest (Fig. 2) [10].

*Acrocomia aculeata* is increasingly recognized as a promising minor crop, offering a diverse array of emerging high-value commodities. These include products derived from its fruits, kernels, nut and leaves, as well as high-content oils suitable for various applications including biofuels, human and animal consumption (vegetable oil and flours), cosmetics, and pharmaceuticals. Furthermore, *A. aculeata* holds significant promise for sustainable biomass production, raising important considerations regarding land use, biodiversity conservation, carbon sequestration and storage, and its potential contributions to climate and environmental protection [2,4]. Biofuels derived from *Acrocomia* could facilitate the transition from fossil fuels to more sustainable energy options in both developed and developing countries. The *Acrocomia* ability to thrive in marginal areas with poor, arid soils, such as rangeland, bare ground, degraded cropland, or grassland, aligns with multiple United Nations Sustainable Development Goals (SDGs): (1) SDG 7 (Affordable and Clean Energy) by providing a renewable fuel source. (2) SDG 13 (Climate Action) by potentially mitigating carbon emissions and promoting climate-resilient agriculture. (3) SDG 15 (Life on Land) by restoring degraded land and potentially mitigating biodiversity loss.

These three species easily occur in anthropized areas, and their dispersion is favored by human interaction [4]. There is evidence that *Acrocomia* was used by Pre-Columbian civilizations as a ritual and edible plant [16–18]. The oldest archaeological record of its utilization in North America is dated from the Pre-Olmec and Olmec civilizations in the southern Mexican state of Veracruz (4000 BP) [19]. The oldest human interaction in South America is with the Paleo-Indians and other proto-Tupi-Guarani civilization in the Brazilian lowlands dated 10,030 BP. The oldest known dated fossil was described in the northern Brazilian state of Pará (Santarén - 11,200 BP), followed by Argentina (8500 BP), Panama (8040 BP), and Mexico (6750 BP) [1,20]. *Acrocomia aculeata* is considered an incipiently domesticated crop and probably share the same evolutionary pressures as other Neotropical domesticated crops for new ecological niche of cultivation and adaptation [18]. Despite the growing economic interest in some *Acrocomia* species, understanding of their genomic information remains limited, especially adaptation strategies and gene flow relationships.

The development of new crop varieties with improved traits such as disease resistance, drought tolerance, and high oil yield can be achieved through the identification of local varieties (ecotypes), genetic markers, and quantitative trait loci (QTL) mapping [21,22]. Understanding the genetic diversity and QTLs of *Acrocomia* palms is essential for identifying desirable characteristics and selecting suitable breeding materials. The ability of crop species to adapt and maintain resilience in response to changing environmental conditions is significantly influenced by genetic diversity [23]. Using loci under selection, breeders can identify and characterize candidate genes responsible for key adaptive traits [24]. Furthermore, research can provide insights on the mechanisms underlying palm diversification in the Neotropic by identifying important evolutionary processes such speciation, adaptation, and dispersal.

This study assessed selective signatures and genome-wide diversity in natural populations of *Acrocomia aculeata*, *Acrocomia totai*, and *Acrocomia intumescens* from different Neotropic local biomes. We used double-digest genotyping by sequencing (GBS) to identify single nucleotide polymorphisms (SNPs) for putative signatures of selection. Our analysis explores the potential role of these SNPs in adaptation, diversification processes, and genetic population structures, particularly in relation to cultivation in diverse environmental contexts, with a focus on important crop traits. This research aims to generate novel insights into the evolution of *Acrocomia* across the Americas and contribute to the domestication and breeding of this promising tropical palm for sustainable vegetable oil production, bioeconomy, and biofuels.

## Results

### SNP discovery in Acrocomia sequencing

Sequencing of the two ddGBS libraries for *A. aculeata* and *A. totai* generated a total of 219,264,253 reads, while the two *A. intumescens* ddGBS libraries generated a total of 554,630,990 reads. After quality-control filtering, the number of retained reads were: 60,786,924 (mean = 779,319.5 reads per sample, SD ± 449,982) for *A. aculeata*; 29,283,574 (mean = 697,227.9 reads per sample, SD ± 355,813) for *A. totai*; and 276,741,447 (mean = 1,990,945.6 reads per sample, SD ± 1,333,058) for *A. intumescens*. After the analysis in Stacks, the final data set had 1997 SNPs for the 78 *A. aculeata* samples (mean depth per locus = 18.7X, SD ± 6.2, 8.8% of missing data). For the 40 *A. totai* samples, 1629 SNPs were identified (mean depth per locus = 17.2X, SD ± 6.4, 2.9% of missing data). For the 131 *A. intumescens* samples a total of 3466 SNPs were identified (mean depth per locus = 11.5X, SD ± 4.4, 13.4% of missing data).

### Putative signatures of selection and population structuration in Acrocomia

Because pcadapt and LFMM are methods that account for the global genetic structure of the data, some interesting patterns may be considered for the analyses of putative signatures of selection. Both analyses suggested that the genetic structure within *A. aculeata* and *A. intumescens* is not explain by the local biome groups assumed according to the classification proposed by Freitas *et al*., [12] but rather the outcome of genomic isolation of biogeographic barriers and historical events. In the case of *A. totai*, a genetic structuring was observed in accordance with local biomes and macro ecoregions (Fig. 3, S1 and S3) suggesting local ecosystem adaptation. Similar DAPC results were obtained, indicating a significant degree of divergence between natural populations from specific biome. The major genetic structure in *A. aculeata* is observed among sampling locations (or biogeographic groups) within and outside Brazilian lowlands (<1500 masl), resulting in the separation of the samples into two clusters along the first principal component (43.88% of the variance), with a distinct divergence between samples from Mexico and Central America and those from Brazil (Fig. 3 and S1). In *A. totai*, samples from Pantanal (PAN) were the most divergent, the Argentinean-Paraguayan Chaco and Espinal (CHE) samples tend to cluster in the same genetic group, while samples from the Brazilian Cerrado (CER) were more admixed than those from the other groups (Fig. 3 and S1). The major genetic divergence in *A. intumescens* was observed between the Brazilian state of Ceará, indicating interrupted gene flow likely due to genetic isolation or a biogeographic barrier, rather than between biomes (Fig. 3 and S1).

**Fig 3.**
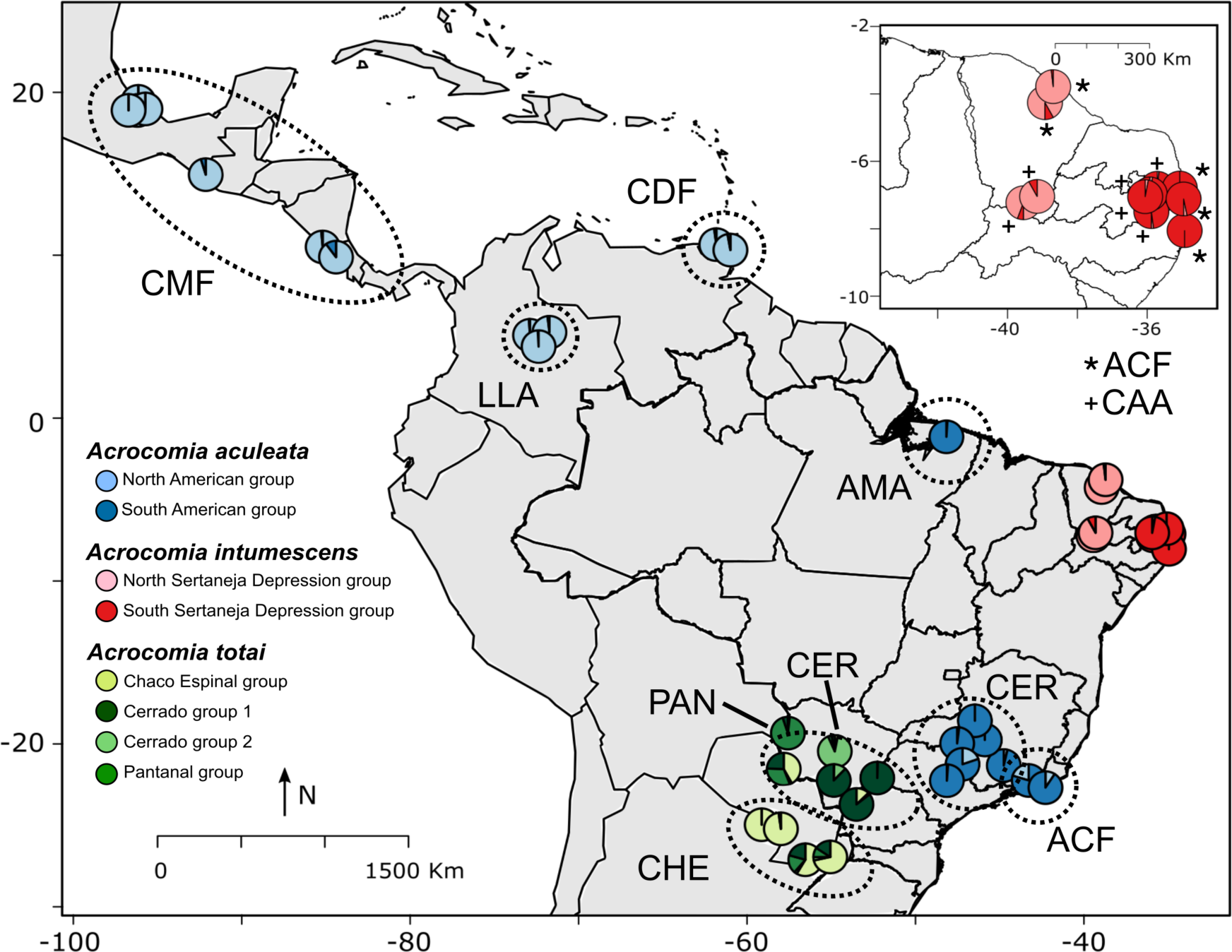
Map of sampling locations for *Acrocomia aculeata* (blue), *A. totai* (green), and *A. intumescens* (red), along with major genetic divergences among local biome groups as suggested by sparse non-negative matrix factorization (sNMF) analyses. Pie charts show the average sNMF ancestry coefficients across the genetic clusters represented by different color shades. Some pie charts were slightly moved to ease visualization. Acronyms follow Table 2 local biomes: CMF Central American Moist Forest; LLA Llanos; CDF Caribbean Dry Forest; ACF Atlantic Coastal Forest; CAA Caatinga; AMA Amazon; CER Cerrado; PAN Pantanal; CHE Chaco and Espinal.

The numbers of outlier SNPs detected for *A. aculeata* were 500 based on pcadapt and 129 based on FstHet, while LFMM indicated 525 loci associated with at least one bioclimatic variable. Of these, 326 markers were considered as putative under selection because they were indicated by at least two of these methods (Fig. 4A). A total of 67 *A. aculeata* sequences with an outlier marker had at least one blastx hit, and 54 had at least one GO annotation. These GO terms could be summarized in 33 different classes (Fig. 5A). For *A. totai*, pcadapt indicated 218 outlier SNPs, FstHet 306, and 446 SNPs were associated to a bioclimatic variable in LFMM. Of these, 192 SNPs were considered as putative under selection (Fig. 4B). A total of 38 *A. totai* sequences with an outlier marker had at least one blastx hit, and 33 had at least one GO annotation. These GO terms could be summarized in 28 different classes (Fig. 5B). For *A. intumescens*, pcadapt indicated 688 outlier SNPs, FstHet 157, and 1024 markers were associated with a bioclimatic variable in LFMM. Of these, 104 SNPs were considered as markers putatively under selection (Fig. 4C). Only 8 *A. intumescens* sequences with an outlier marker had at least one blastx hit, and 2 had GO annotations. These GO terms could be summarized in 7 different classes (Fig. 5C). For the three species, the most common GO annotations were related with the biological processes of “metabolic process” and “cellular process”, and the molecular functions of “binding” and “catalytic activity”. GO annotations related to “response to stimulus” and “regulation” were also frequent for the proteins with similarities to the sequences with outlier SNPs of *A. aculeata* and *A. totai*.

**Fig 4.**
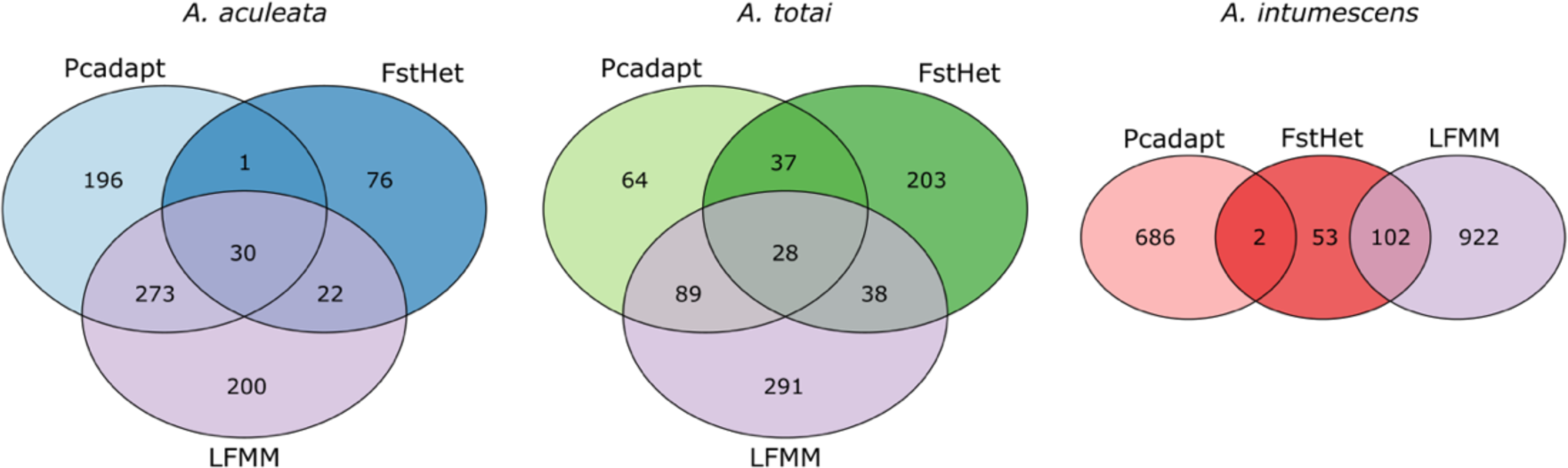
Venn diagrams showing the number of loci identified as outliers with different methods. The outlier SNPs detected in Pcadapt are based on the genetic structure of the PCA, while in FstHet, they are based on the betahat statistic. In LFMM, the outliers are associated with bioclimatic variables. A) *Acrocomia aculeata*, B) *A. totai*, and C) *A.intumescens*.

**Fig 5.**
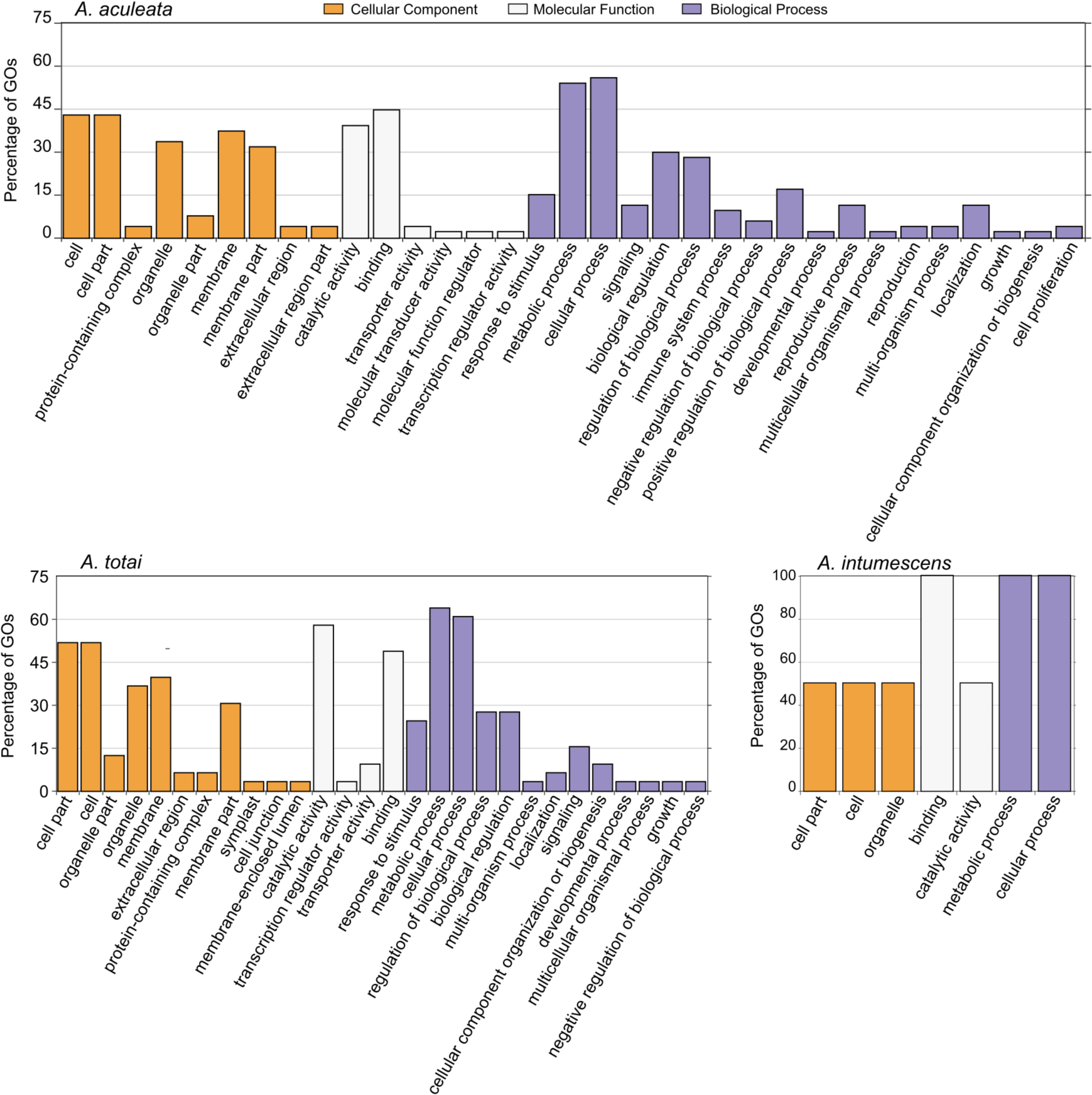
Summary of Gene Ontology (GO) annotations associated with proteins. The outlier SNPs in gene sequences associated with proteins are deposited in GenBank. GO terms summarize annotations according to cellular components, molecular functions, and biological processes. A) *Acrocomia aculeata*, B) *A. totai*, and C) *A.intumescens*.

These genes exhibited high similarity to genes annotated and described in palm species such as *Elaeis guineensis*, *Phoenix dactylifera* L., and *Cocos nucifera* L., all of which have fully sequenced genomes (Table 1, S2). The genes observed were related to many functions, including oil production metabolism/catabolism, carotenoid biosynthesis, plant growth and development, organ sizes, root, flowering, pathogen resistance, and biotic and abiotic stresses. The most relevant loci identified as outliers in different methods and for each species are listed in Table 1.

**Table 1.** Most relevant gene loci with putative selective signatures in *Acrocomia aculeata*, *A. totai*, and *A. intumescens*.

| *Acrocomia aculeata* |  |  |  |  |  |  |
| --- | --- | --- | --- | --- | --- | --- |
| Sequence name | Sequence desc. | Species | Function | E-Value | Similarity | Reference |
| CLocus_16512 | Carotenoid cleavage dioxygenase 7 (chloroplastic) | *Elaeis guineensis* | Carotene catabolic process, Strigolactone biosynthesis that modulates plant growth, reproduction, senescence | 4.87E-09 | 89.7 | [25–28] |
| CLocus_27100 | Putative WRKY transcription factor 19 | *Elaeis guineensis* | Modifies the content of fatty acids and influences the accumulation of seed oil; Abiotic stress responses, and plant pathogen interactions | 1.96E-10 | 100.0 | [29,30] |
| CLocus_25725 | Lipid binding protein | *Elaeis guineensis* | Lipid biosynthesis Lipid transport; Lipid binding | 2.37E-09 | 100.0 | [31] |
| CLocus_317 | Polyprenol reductase 1 isoform X1 | *Elaeis guineensis* | Abiotic stress responses, and plant pathogen interactions, lipase activity | 9.80E-08 | 96.6 | [32,33] |
| CLocus_11291 | Patatin-like protein 2 | *Cocos nucifera* | Lipid Metabolism (phospholipase activity; lipid catabolic process; acylglycerol lipase activity); Resistance to pathogens | 3.83E-11 | 96.7 | [34,35] |
| CLocus_11820 | carotenoid cleavage dioxygenase 7, chloroplastic | *Phoenix dactylifera* | Carotene catabolic process, modulates plant growth, reproduction, senescence | 4.37E-09 | 90.0 | [25,28,36] |
| CLocus_24655 | pentatricopeptide repeat-containing protein At5g15340, mitochondrial | *Elaeis guineensis* | Zinc ion binding. Influence organellar expression function and, consequently, on photosynthesis, respiration, plant development, and environmental responses | 9.48E-10 | 100.0 | [37] |
| CLocus_12388 | pentatricopeptide repeat-containing protein, mitochondrial-like | *Cocos nucifera* | Zinc ion binding. Influence organellar expression function and, environmental responses | 1.20E-06 | 86.7 | [37] |
| CLocus_16644 | Polyprenol reductase 2 | *Phoenix dactylifera* | Essential in dolichol biosynthesis | 7.80E-10 | 100.0 | [32,33] |
| CLocus_17318 | Carotenoid cleavage dioxygenase 8 homolog B (chloroplastic) | *Morus notabilis* | Apocarotenoids, carotenoid cleavage dioxygenases (CCDs), inhibit shoot branching. Also active on other carotenoid substrates like lycopene or zeaxanthin | 1.92E-11 | 100.0 | [38,39] |
| CLocus_18527 | Protein OBERON 3 | *Phoenix dactylifera* | Embryonic root meristem initiation, rooting | 2.16E-10 | 100.0 | [40,41] |
| CLocus_21286 | GRF-interacting factor 1 | *Elaeis guineensis* | Leaf development and growth, reproductive system development and growth | 3.67E-09 | 96.6 | [42,43] |
| CLocus_22454 | Momilactone A synthase | *Cocos nucifera* | Momilactone A biosynthesis, diterpenoid secondary metabolites, involved in the defense mechanism of the plant produced in response to attacks | 9.13E-12 | 100.0 | [44,45] |
| CLocus_23382 | Phosphoinositide phospholipase C 2 | *Elaeis guineensis* | Signal transduction in events such as guard cell signaling, salt stress, osmotic stress, acquired resistance, Nod factor signaling, drought, systemic acquired resistance, carbon fixation in C4 plants, and response to the pathogen | 4.27E-12 | 100.0 | [46] |
| CLocus_29621 | BTB/POZ domain and ankyrin repeat-containing protein NPR2 | *Cocos nucifera* | Plant defense and plant disease resistance | 4.59E-10 | 100.0 | [47] |
| CLocus_30624 | Subtilisin-like protease SBT1.5 | *Elaeis guineensis* | Plant-pathogen recognition and immune priming | 5.86E-09 | 96.6 | [48] |
| CLocus_35203 | BTB/POZ domain and ankyrin repeat-containing protein NPR2 | *Elaeis guineensis* | Plant defense and plant disease resistance | 3.68E-10 | 100.0 | [47] |
| CLocus_21703 | FRIGIDA-like protein 4a | *Elaeis guineensis* | Regulates flowering induction, flowering time | 2.22E-04 | 100.0 | [49] |
| CLocus_25856 | Putative pentatricopeptide repeat-containing protein At5g37570 | *Elaeis guineensis* | Organelle biogenesis and function, photosynthesis, respiration, plant development, and environmental responses | 1.15E-10 | 96.6 | [37] |
| CLocus_32860 | ABC transporter D family member 1 | *Capsicum annum* | Lipid metabolism (long-chain fatty acid transporter activity, Long-chain fatty acid import into peroxisome; fatty-acyl-CoA transport), Organ growth, plant nutrition, plant development, response to abiotic stress, and the interaction of the plant with its environment | 2.37E-04 | 100.0 | [35,50] |
| CLocus_82942 | Endoglucanase 8-like | *Elaeis guineensis* | Synthesis, remodeling and turnover of cell wall components in symbiotic and hostile plant-microbe interactions | 1.09E-06 | 100.0 | [51] |

| *Acrocomia totai* |  |  |  |  |  |  |
| --- | --- | --- | --- | --- | --- | --- |
| Sequence name | Sequence desc. | Species | Function | E-Value | Similarity | Reference |
| CLocus_17145 | GDSL esterase/lipase | *Elaeis guineensis* | Lipid metabolism: produce and storage oil in seeds, cuticular lipids to cover and decorate organ surfaces, oxylipins, and other signaling molecules | 7.53E-11 | 100.0 | [52,53] |
| CLocus_37219 | Subtilisin-like protease SBT1.2 | *Elaeis guineensis* | Plant-pathogen recognition and immune priming | 3.18E-11 | 100.0 | [48] |
| CLocus_15942 | Low affinity sulfate transporter 3 | *Cocos nucifera* | Plant response to drought and salinity stress | 7.41E-10 | 100.0 | [54] |
| CLocus_16267 | Mitogen-activated protein kinase kinase kinase NPK1 | *Elaeis guineensis* | Regulates innate immunity and development in plants | 1.33E-08 | 93.1 | [55] |
| CLocus_20717 | Fatty acid desaturase 4, chloroplastic | *Cocos nucifera* | Influence levels of susceptibility to multiple stresses, including insect infestations | 2.21E-09 | 100.0 | [56] |
| CLocus_21866 | Phosphatidylinositol 4-kinase alpha 1 isoform X2 | *Elaeis guineensis* | Essential for pollen, embryonic, and post-embryonic development also cell signaling | 5.17E-10 | 100.0 | [57] |
| CLocus_24114 | 9-cis-epoxycarotenoid dioxygenase NCED6, chloroplastic | *Cocos nucifera* | Increases drought tolerance, multi-abiotic stress tolerance, regulates plant growth, also enhances seed dormancy | 3.42E-13 | 96.6 | [58–60] |
| CLocus_26989 | Momilactone A synthase | *Cocos nucifera* | Drought, salinity, and oxidative stress conditions tolerance | 1.58E-11 | 100.0 | [61,62] |
| CLocus_29376 | Subtilisin-like protease SBT1.2 | *Elaeis guineensis* | Plant-pathogen recognition and immune priming | 2.36E-10 | 93.3 | [48] |
| CLocus_37352 | Protein OBERON 3 | *Phoenix dactylifera* | Embryonic root meristem initiation, rooting | 2.21E-10 | 100.0 | [40,41] |
| CLocus_54281 | Phospholipase A2-alpha | *Elaeis guineensis* | Signaling roles during plant abiotic and biotic stress responses | 2.52E-09 | 96.6 | [63] |
| CLocus_12178 | Protein ENHANCED DISEASE RESISTANCE 4 | *Cocos nucifera* | Regulation of defense response to fungus | 1.61E-05 | 93.3 | [64] |

| *Acrocomia intumescens* |  |  |  |  |  |  |
| --- | --- | --- | --- | --- | --- | --- |
| Sequence name | Sequence desc. | Species | Function | E-Value | Similarity | Reference |
| CLocus_111127 | Phosphatase 2C BIPP2C1 | *Elaeis guineensis* | Plant signal transduction processes and stress signaling | 3.23E-09 | 100.0 | [65] |

## Discussion

The focus of population genetic research has rapidly shifted from spatially neutral genetic processes to adaptive genetic processes, driven by advancements in SNP genotyping techniques for studying crop traits and domestication processes. Considerable deviation of outlier markers from the distribution of a specific statistical evaluation under a particular model can serve as the basis for methods aimed at detecting selective signatures for understanding adaptation events [24,66]. A significant future development in sustainable agriculture will involve the domestication of novel native plant species, such as *Acrocomia*, to facilitate the transition towards cleaner energy sources. Below, we discuss the most interesting findings regarding outlier SNPs in the adaptation process of *Acrocomia*, their biological and evolutionary significance, and their implications for breeding efforts.

### The adaptation and evolutionary process in *Acrocomia*: Dispersal and Biogeographic Barriers

*Acrocomia* phenotype has been shaped by ecological interactions that have influenced the evolution of specific traits for its survival [67]. *Acrocomia* species are distributed across tropical America, extending from North America (Northern Mexico) to South America, inhabiting areas transitioning from tropical and subtropical rainforests (moist broadleaf forests) to savannah and xeric shrubland regions, including dry broadleaf forests (semi-arid and arid ecosystems) (Fig. 2). Their wide distribution suggests their ability to adapt to various soil and climate conditions. sNMF analyses can help identify genetic signatures associated with adaptation to specific environmental conditions or phenotypic traits. Our findings provide insights into the genetic structure and the association of specific SNP loci under selection forces in the adaptation process of *Acrocomia aculeata*, *A. totai*, and *A. intumescens* (Fig. 4, Table 1).

Our results support the division of *A. aculeata* into two major gene pools: a North American group and a South American group, consistent with previous observations based only on neutral loci [3] (Fig. 2 and 3). Notably, we found no significant differences in genetic structure within minor local biomes within each gene pool, indicating that the Amazon basin likely served as a source region for diversification across the species distribution. This pattern in genetic structure is also present in other crop species such as common beans and maize [68,69]. Both dispersal and biogeographic barriers likely influenced the genomic structure of *Acrocomia aculeata*. Also, *A. aculeata* exhibited the greatest number of putative selective signatures (Table 1; S1 and S2) in genes associated with fatty acid and triacylglycerol biosynthesis, carotenoid biosynthesis, pathogen resistance and defense, as well as genes specialized in adapting to abiotic and environmental stress. From the nine species of the genus, *A. aculeata* exhibits the highest fruit pulp oil content (65% w/w dry basis) [4].

Individual plant fitness is influenced by animal–plant interactions, encompassing mutualisms such as seed dispersal and antagonistic relationships like herbivory and seed predation [67]. For example, the morphology of *Acrocomia aculeata* fruit, including its size, color, and content, appears to have undergone correlated evolution as predicted by the seed-dispersal syndrome hypothesis before the beginning of its domestication process through very recently human interaction (∼10,000 BP) [1,20]. Additionally, spines and other mechanical defense features are common mechanisms employed by plants to avoid herbivory. Numerous studies on palms indicate that spinescence reduces herbivory by large mammals [70]. The same herbivory and seed dispersal pattern is present in A. *totai* and *A. intumescens*.

Frugivorous mammals have a preference for consuming dull-colored fruits, such as those that are green, yellow, brown, or pale orange [67,71] (Fig. 1). It is hypothesized that the coccoid type fruit (drupe) of *Acrocomia* ancestor species may have been consumed by large herbivores in the past, with migration events like the Great American Biotic Interchange (GABI; 2.5 million BP) likely influencing its distribution through seed dispersal [71]. Consequently, certain observable fruit traits and defenses against herbivory, such as spiny trunks and hard epicarp, could be considered anachronic, representing interactions that occurred in the past [71]. The term “anachronic” denotes traits or features that are outdated or no longer relevant in the current context but may have been adaptive or functional in the past [67,71]. While most large mammals are now extinct in the Neotropics following the Late Quaternary Extinction episode, Neotropical palms like *Acrocomia* diversified in an ecological environment where mammalian assemblages were abundant in large-bodied species, approximately from 2.6 million to 23,000 BP [67,72,73]. However, our findings suggest only a certain level of connectivity between populations within the two major gene pools. Currently, cattle and humans are regarded as the primary contributors to seed dispersal for this species, highlighting its recent close association with human activity [70,74].

Archaeological evidence encompassing Pre-Columbian civilizations across the neotropics, from Mesoamerica (northern Mexico, Central America) to South America, suggests the cultivation and ritualistic utilization of *Acrocomia aculeata* [19,20]. The presence of *Acrocomia* in these regions indicates a potential influence of Amerindian migration on species diversification and offers insights into its domestication process. Such historical records reinforce the relationship between indigenous populations and *Acrocomia*, potentially shaping its genetic diversity and evolutionary trajectory through intentional cultivation and cultural practices. Further investigation is necessary to elucidate the specific historical events and ecological factors shaping the observed genetic structure and center of origin.

*Acrocomia totai* exhibits a pronounced genetic structuring, showing a clear differentiation between populations from the Brazilian Cerrado, Pantanal, and Chaco and Espinal (Fig. 2 and 3). This contrasts with previous studies focused only on neutral loci using SNPs and microsatellites where they observed less genetic structuring among populations [3,75]. This indicates that the populations of *A. totai* are experiencing a recent diversification and ongoing speciation processes.

*A. totai* has a subtropical distribution and is adapted to temperate conditions, with selective signatures found in genes associated with fatty acid and triacylglycerol biosynthesis, pathogen resistance and defense, as well as genes specialized in adapting to abiotic and environmental stress (Table 1; S1 and S2). Pantanal populations exhibit lower levels of admixture compared to other genomic groups within *A. totai*, particularly those distributed in the Brazilian Cerrado. The Cerrado lowlands and ecotones with other local biomes serve as a bridge, including for the South American gene pool of *A. aculeata*. Hybridization between these two species has been reported [3]. Phenotypic differences were also observed in other studies, compared to fruits from the Pantanal, fruits from Cerrado had a higher proportion of epicarp and a lower proportion of kernels [5]. The Pantanal biome, an active sedimentary basin characterized by faults and earthquakes, experiences subsidence and depressions that are highly susceptible to flooding. During the late Pleistocene (14,000–10,000 BP), arid conditions predominated, possibly allowing *A. totai* to recently colonize the biome [76]. The Pantanal lacks endemic tree species, with the majority of terrestrial species being immigrants from the Cerrado [77]. Similar biogeographic patterns are observed in the palm *Copernicia alba* Morong. [78]. The domestication process of *A. totai* remains poorly understood, although archaeological records indicate its use for fiber production in Argentina [20]. Human utilization varies among *Acrocomia* species, with *A. totai* being utilized for the yellowish pulp of the fruit for flour production, and the oil derived from the seed, which are the two products of greatest interest [5,14,79].

*A. intumescens* presented a genetic structure influenced by an allopatric isolation across various phytophysiognomies within Caatinga, rather than by variations in biomes (Fig. 2 and 3) [80,81]. Specifically, notable genetic structuring was observed between populations in the North Sertaneja depression and those transitioning from the Borborema highlands to the South Sertaneja depression within the Caatinga biome. The Caatinga exhibits high levels of endemism, with species adapted to survive in its arid climate characterized by drought conditions. The long-term stability of the Caatinga, along with the assembly of ancient plants, has been significantly influenced by aridification processes, while recent vegetation shifts and climate change have driven in situ diversifications. The increased environmental variability has led to the appearance of modern species through Pleistocene/Pliocene (2.6 million BP) ecological specialization [82]. This could elucidate the phylogenomic proximity observed between *A. aculeata* and *A. intumescens*, suggesting a shared common ancestor in the past and probably, *Acrocomia* recently colonized and diversified in the Caatinga [83]. This shared ancestry may be attributed to the dominance of tropical rainforests in the region, which connected the area to the Amazon during the Early Cenozoic period [3,84]. This pattern of endemism and genetic differentiation is shared with other plant species in the Caatinga, including other palms like *Copernicia prunifera* (Miller) H.E. Moore, legumes (*Coursetia*, *Vatairea*, and *Luetzelburgia* genera), as well as other clades such as *Conopophaga cearae* (Aves. common name: Caatinga gnateater) and lizard species [81,82,85,86]. In *A. intumescens*, the only selective signature found in a gene was associated with stress signaling.

The close relationships observed among *Acrocomia* species may be attributed to the radiation of the Bactridinae subtribe towards the end of the Eocene (23 million BP), when the ancestor of *Acrocomia* diversified [87]. This period of divergence coincided with the terminal Eocene cooling event, characterized by substantial climatic changes that led to accelerated turnover in flora and fauna [88]. This pattern is also evident in the conserved genomic structures and similarity of *Acrocomia aculeata*, *A. totai* and, *A. intumescens* in their plastomes architecture and nuclear gene phylogeny, low pairwise Fst indicating a close relationships [3,84]. The high admixture between populations and the short branch lengths observed in phylogenomic studies, which separate the three species, presents challenges in establishing unambiguous intergeneric relationships within the genus *Acrocomia*. This may suggest a large-scale species-level extinction followed by rapid diversification of surviving lineages.

Correlations between allele frequencies and climatic variables in the LFMM analysis revealed five climatic variables influencing adaptation in the three species: annual mean temperature (BIO1), mean diurnal range (BIO2), isothermality (BIO3), mean diurnal range (BIO12), and precipitation of the driest month (BIO14), indicating selection as a driving force of evolutionary adaptation (Fig. S2). Similar patterns were observed in photosynthesis rates and fruit pulp mass development in other *Acrocomia* studies and in the neotropical palm *Mauritia flexuosa* Mart., which is influenced by BIO1, BIO2, and BIO14 [7,89]. Although *Acrocomia* can thrive in diverse climatic conditions occurring from humid to semi-arid ecosystems, they are exclusively found in open areas, often influenced by human activity. Our results are supported by previous studies that suggest temperature played a significant role in the evolution of the climate niche in Neotropical palm species [9,90].

The sNMF analysis (Fig. S2) showed low differentiation among populations from different biomes in putative SNPs under selection. However, there was evidence of admixture in *Acrocomia totai*, particularly between natural populations from Cerrado in Paraguay, Mato Grosso do Sul (Brazil), Parana (Brazil), and Sao Paulo (Brazil). Biogeographic isolation by barriers is an important driver of genetic differentiation in palm as described by Eiserhardt *et al.:* “The significant effect of spatial distance on clade turnover in American palms can thus be interpreted as evidence for dispersal limitation on evolutionary timescales caused either by barriers or by time” [9]. Even though Eiserharth’s results focus on macroevolutionary diversification and phylogenetic turnover by dispersal and niche evolution, they indicate that barrier distance explains more variation in diversity than spatial distance. We observe a similar pattern in our results, where biogeographic barriers are more relevant in shaping genetic structure than biomes proximity and microclimatic variables in *Acrocomia* niches (Fig. 3).

Palms are not naturally adapted to withstand extremely high or low temperatures. Interestingly, the evolution of these geographic range restrictions does not appear to be a limiting factor for *Acrocomia*, despite the fact that water supply most likely limits the regional ranges of many Neotropical palms [91]. *Acrocomia* appears to be highly adapted to semi-arid ecosystems. The distributions of Neotropical palm species are assumed to be impacted by edaphic environments, despite the absence of empirical data. It is known that the natural occurrence in Brazilian populations of *Acrocomia aculeata* is associated with eutrophic (high fertility) soils with medium to clayey textures and an average pH of 5.5 [92]. However, further investigation is needed for the North American gene pool characteristics.

Domestication syndrome in palms is poorly understood [17,93]. Detecting selective sweep regions associated with domestication in *Acrocomia* is crucial. These genes probably have undergone strong positive selection during this process. Selective sweeps are characterized by a reduction in genetic diversity around the selected allele due to the rapid increase in frequency of advantageous alleles favored by human-driven selection [94]. Maybe there is a heterogeneous origin in the domestication of *Acrocomia*. However, the existence of several founder lineages and the link between a mosaic ancestry patterns only will be clear with a sequenced reference genome for the three species. The identification of loci under selection can be greatly enhanced by having access to a reference genome [22]. *Acrocomia* may respond polygenically to environmental variation like other palms [35,89,95], meaning that selection may result in minimal changes in allele frequencies, leading to a poor adaptation signal at a single locus.

Our results demonstrate that genetic structure in populations from different biomes is homogenized within the same biogeographic regions due to gene flow. This also suggests the potential for migration to disperse advantageous alleles among populations sharing the same biogeographic region, leading to allele fixation in genes involved in adaptation to environmental change. This potentially explains *Acrocomia*’s tolerance to seasonality and sheds light on the species’ interaction with humans and its domestication process. We suggest investigating the functionality of distinct biological pathways under diverse conditions using a transcriptomic approach for understanding better its domestication syndrome and its potential resilience to climate change [7,90]. Also, conservation strategies can be formulated by identifying genetically distinct populations that may require special attention for conservation.

### Genes putatively involved in adaptation and implications for domestication in ***Acrocomia***

A high-quality reference genome sequence and annotation at chromosomal level of plant species is essential for genetic research on crop breeding and domestication [21]. *Acrocomia* is considered an incipiently domesticated plant and does not have a reference genome yet, so a population genomics approach using GBS or RAD-Seq can help investigate the evolutionary mechanisms underlying its diversification and variation. Significant genomic alterations have occurred during plant domestication as a result of evolutionary processes such as genetic drift and artificial selection. The study of artificial selection has used genomic techniques for detecting signatures putatively under selection, which focus on identifying polymorphisms in trait loci that have been disguised by neutral variation in species under a domestication process [22].

Despite lacking a complete reference genome for *Acrocomia*, we were able to analyze genes potentially under selection. To do this, Blast2GO and Blastx use different reference genomes of related palm species to compare the loci where signatures of potential adaptation were located. The most common similarity matches between our GBS tags and annotated sequences were found mostly in homologous genes of *Elaeis guineensis* (oil palm), *Cocos nucifera* (coconut), and *Phoenix dactylifera* (date palm) (Table 1). For example, phylogenomic studies utilizing chloroplast genome sequences of *Acrocomia aculeata* and *Elaeis guineensis* demonstrated that all nodes had a posterior probability of 1.0 (PP = 1.0), indicating a close relationship between these species [84]. Here, we present some relevant candidate genes that exhibit selection signatures and could be involved in the adaptation processes (and maybe the domestication) of *Acrocomia*.

Some significant selection signatures were identified in genes associated with fatty acid and triacylglycerol biosynthesis pathways in *Acrocomia aculeata* (Table 1). For example, loci such as CLocus_25725 correspond to the gene encoding a lipid-binding protein that facilitates lipid transport. CLocus_11291 corresponds to the gene encoding a patatin-like protein 2, which exhibits phospholipase activity in lipid catabolic processes and acylglycerol lipase activity. Additionally, CLocus_32860 was found in the gene encoding an ABC transporter D family member 1, involved in lipid metabolism functions such as the transport of long-chain fatty acids, import of long-chain fatty acids into peroxisomes, and fatty-acyl-CoA transport. Furthermore, CLocus_27100 corresponds to the gene encoding a putative WRKY transcription factor that modulates the content of fatty acids and influences the accumulation of seed oil. Additionally, we found signatures like CLocus_24655 and CLocus_12388 related to genes involved in zinc ion binding processes. In the research led by Couto *et al.*, candidate genes related to oil production traits in *A. aculeata* were identified [96]. The genome-wide association study (GWAS) suggested that candidate genes controlling oil production were associated with metal ion binding and correlated with traits such as fruit oil content, fruit pulp fresh mass, leaf number, and leaf length. In *Acrocomia totai*, the locus CLocus_17145 was present in the gene encoding GDSL esterase/lipase, which mobilizes the lipids stored in seeds and plays a major role in seed germination and early seedling establishment [53]. These SNPs may be of agronomic importance because one of the primary objectives of the *Acrocomia* breeding program is to increase oil content for industrial purposes such as biofuels.

These fatty acids and triacylglycerols biosynthesis pathways were also reported in the fine mapping and cross-validation of QTLs linked to fatty acid composition in different varieties of *Elaeis guineensis* made by Ding *et al.* [53]. The Blast results for their QTL included genes and transcription factors linked to diacylglycerol acyltransferase (DGAT1) and long chain acyl-CoA synthetase. The synthesis of triacylglycerol and fatty acids occurs in different compartments within plants. Specifically, new fatty acid synthesis takes place in the plastid (leucoplast), where the acyl carrier protein (ACP) holds the fatty acid chain as it elongates. Then, acyl-ACP thioesterases hydrolyze the acyl-ACPs, releasing non-esterified fatty acids. These fatty acids are then exported to the endoplasmic reticulum, where they assemble to form triacylglycerol [35,97].

Another significant finding were signatures associated in carotenoid biosynthesis pathways (Table 1). In *Acrocomia aculeata*, loci such as CLocus_16512 and CLocus_11820 were associated with genes encoding carotenoid cleavage dioxygenase 7 and carotenoid cleavage dioxygenase 8 homolog B, which play roles in carotene catabolic processes and the production of apocarotenoids. These enzymes, known as carotenoid cleavage dioxygenases (CCDs), also influence shoot and branching inhibition and act on various carotenoid substrates like lycopene or zeaxanthin. Additionally, in *A. totai*, the locus CLocus_24114 was found in the gene encoding 9-cis-epoxycarotenoid dioxygenase NCED6, which is involved in regulating plant growth and seed dormancy. Carotenoids are vital for plant growth and development as they serve as precursors for the synthesis of plant hormones such as strigolactones and abscisic acid (ABA) [26,27]. From a nutritional perspective, the high content of carotenoids in crops is an attractive trait, offering benefits such as mitigating vitamin A deficiency through dietary intake of plant-derived carotenoids. For example, the increasing popularity of orange carrots may be attributed, at least in part, to the numerous studies highlighting the health advantages associated with carotenoids [98].

Additionally, we observe the presence of outlier SNPs in putative resistance genes from different classes, indicating that natural populations under selection possess important genetic resources for crop defense against pathogens (Table 1). For instance, outlier SNPs such as CLocus_27100, CLocus_317, CLocus_22454, CLocus_29621, CLocus_35203, and CLocus_11291 were identified in genes involved in plant defense and disease resistance responses in *Acrocomia aculeata*. In *A. totai*, SNPs such as CLocus_37219, CLocus_29376, and CLocus_12178 were associated with genes involved in plant-pathogen recognition, immune priming, and regulation of defense response to fungus, respectively. Currently, there is no evidence of pathogens, pests, or diseases in either natural populations or commercial plantations of *A. aculeata* that are currently in the production phase located in Brazil’s central region [4]. Phytosanitary issues with this novel crop would likely arise during the domestication bottleneck in response to genetic diversity and loss of heterozygosity.

We have identified signatures in genes specialized in adapting to abiotic and environmental stress in *Acrocomia aculeata*, such as CLocus_317, CLocus_12388, CLocus_23382, CLocus_25856, and CLocus_32860 (Table 1). In *Acrocomia totai*, a signature like CLocus_15942 and CLocus_24114 were found in the gene encoding low-affinity sulfate transporter 3 and 9-cis-epoxycarotenoid dioxygenase NCED6 respectively, responsible for the plant’s response to drought and salinity stress [54]. Similarly, in *Acrocomia intumescens*, the only signature (CLocus_111127) present on a gene was also associated with stress responses. This may help elucidate how *Acrocomia* species can withstand extreme temperatures and droughts. Drought tolerance is a polygenic, complex quantitative trait [99]. Developments in agricultural physiology and genetics have contributed with important insights into drought tolerance in palm species [95,99,100]. Therefore, improving yield under water-deficient conditions is a major objective in plant breeding. Increasing drought tolerance through traditional breeding is a slow process [101]. *Acrocomia*’s inherited characteristics give it the advantage of being adapted to semi-arid, even arid regions like the Brazilian Caatinga. For the majority of non-model species, the mechanisms driving drought tolerance remain poorly understood and vary among species. In comparison to domesticated species, incipiently or less domesticated populations may have a larger range of ecological adaptations [23]. Crop adaptability to perturbed areas depends on its resistance and resilience to environmental stress, which could be crucial for *Acrocomia*’s adaptation in the face of climate change.

### Challenges of reduced representation sequencing (GBS/RAD-seq) in adaptation studies

Reduced representation sequencing methods in adaptation studies such as genotyping-by-sequencing (GBS) or restriction-site associated DNA sequencing (RAD-seq) employ high-throughput sequencing to generate high genome-wide marker data. The advantages of these techniques is their capacity to develop genotyping assays without requiring prior genomic knowledge or substantial costs without needing a reference genome like *Acrocomia* [21]. GBS serves as a great alternative for investigating unconventional food plants, natural populations, and wild relatives. Additionally, it has been employed to assess diversity and gene flow between crops and their wild types, as demonstrated in Brazilian manioc varieties and other crops [24]. Some challenges in adaptation analysis using GBS data include allelic dropout [102], missing targets in resequencing, false positives selection signatures [103] and the management of missing data [104]. However, the identification of false positives may arise from departures from the model’s assumptions and covariance with sampling techniques, population structure, and demographic variables [103]. Generally, combining the SNPs identified by various outlier tests is one approach to address these limitations [24,94].

Meanwhile, it is becoming increasingly evident that many crops have complex ancestries, necessitating additional archaeogenomics data [22]. Because of its wide distribution, to gain a better understanding of the domestication process of *Acrocomia* and possible genes targeted by human selection, a more comprehensive examination of variables including the diversity of wild population substructure and the proportional genetic gains of various natural subpopulations is recommended, especially in the North American genepool. A pangenomic approach is crucial to consider because plant genomes are dynamic, containing a reservoir of genetic diversity that enables adaptation to different biomes [23]. Future crop varieties will likely require a broader range of genes than those provided by single reference genomes. Developing a pangenome of *Acrocomia* could capture the genomic diversity present in the various gene pools observed in our research.

### The promising aspects of *Acrocomia* and future perspectives

After exploring various aspects of population genetics, adaptation processes, and potential domestication in *Acrocomia* species, our study presents a comprehensive understanding of the genetic structure, evolutionary history, and adaptive mechanisms within this genus. Through advanced genomic techniques such as SNP genotyping and adaptive signatures analyses, we explored the relationships between different *Acrocomia* species and their adaptation to diverse environmental conditions across neotropical biomes. A high degree of diversity was observed in *Acrocomia* species from various neotropical biomes. This variation may have arisen from the crop’s natural history, domestication process and its cultivation in diverse ecosystems with varying human preferences. It is plausible that some of the selection signals are associated with desirable agronomic traits, which could be crucial for the breeding of these palms.

*Acrocomia* emerges as a promising crop due to its multipurpose potential, offering prospects in oil production, biofuels, food, and pharmaceutical industry [105]. Notably, the oil extracted from *A. aculeata* exhibits quality comparable to palm oil (*Elaeis guineensis*) while requiring lower water resources and demonstrating adaptability to semi-arid to arid environments and degraded ecosystems. Given that approximately 80% of Brazilian grasslands are classified as degraded, intercropping with perennial plants like *Acrocomia* emerges as a promising solution for combating soil degradation [4,7]. Currently, Brazil is leading its breeding programs through public-private initiatives focusing on *Acrocomia aculeata* and *A. totai* [106]. It is projected that experimental and commercial plantations of *A. aculeata* will reach an area of 200,000 hectares in the next decade [107], with a guaranteed market for the oil and biodiesel production. Significant research progress has been made, particularly in seed germination, seedling production, and integrated agricultural systems, although molecular breeding remains necessary [2,4]. Prioritizing the sequencing and annotation of the entire genomes of the three most economically important species in the genus is crucial for future genomic efforts in their domestication process. Addressing challenges such as genotype-environment interaction, biodiversity prospecting for phenotypic breeding ideotypes (plant models), and crop management are necessary. Most research efforts have focused only on the South American gene pool.

Our findings reveal significant genetic structuring within and between species, highlighting the influence of dispersal, biogeographic barriers, and historical ecological interactions on the evolution of *Acrocomia*. We observed distinct genetic signatures associated with adaptation to specific environmental factors, including climatic variables, pathogen resistance, and stress responses. Notably, *Acrocomia* species exhibit potential resilience to climate change, particularly in semi-arid ecosystems, making them promising candidates for sustainable agriculture and biofuel production. Furthermore, our study sheds light on the potential domestication of *Acrocomia* species, emphasizing the importance of genetic resources and candidate genes associated with desirable agronomic traits such as oil content, carotenoid biosynthesis, and stress tolerance. While facing challenges inherent to reduced representation sequencing methods, such as GBS, our research underscores the necessity of comprehensive genomic resources, including reference genomes and pangenomic approaches, to fully elucidate the domestication syndrome and facilitate breeding efforts.

*Acrocomia* is a valuable crop offering opportunities for future industries while contributing to sustainable development and biodiversity conservation. By prioritizing collaborative research efforts in private-public programs, genetic conservation, and policy support, different stakeholders can unlocked the potential of genetic diversity and adaptive capacity of *Acrocomia* species to address global challenges, including food security, renewable energy production, and climate resilience in agroecosystems.

## Materials and methods

### Plant material and DNA isolation

We sampled leaves from natural populations, 78 for *Acrocomia aculeata*, 40 for *Acrocomia totai*, and 131 for *Acrocomia intumescens*, for a total of 249 samples (Table 2). The collections of these samples were registered in the Brazilian National Council for Genetic Patrimony CGEN (numbers A69E071 and A5D139D). The leaves were dehydrated using silica gel and stored in paper bags at -20 °C. Following Doyle and Doyle’s protocol, we extracted whole genomic DNA from 50 mg leaf samples. Agarose gel electrophoresis (1% w/v) with GelRed stain (Sigma-Aldrich) was used to assess DNA quality and integrity. We quantified and normalized DNA concentrations to 30 ng/μL using the dsDNA BR Assay quantification kit for the Qubit3 fluorometer (Invitrogen).

**Table 2.**
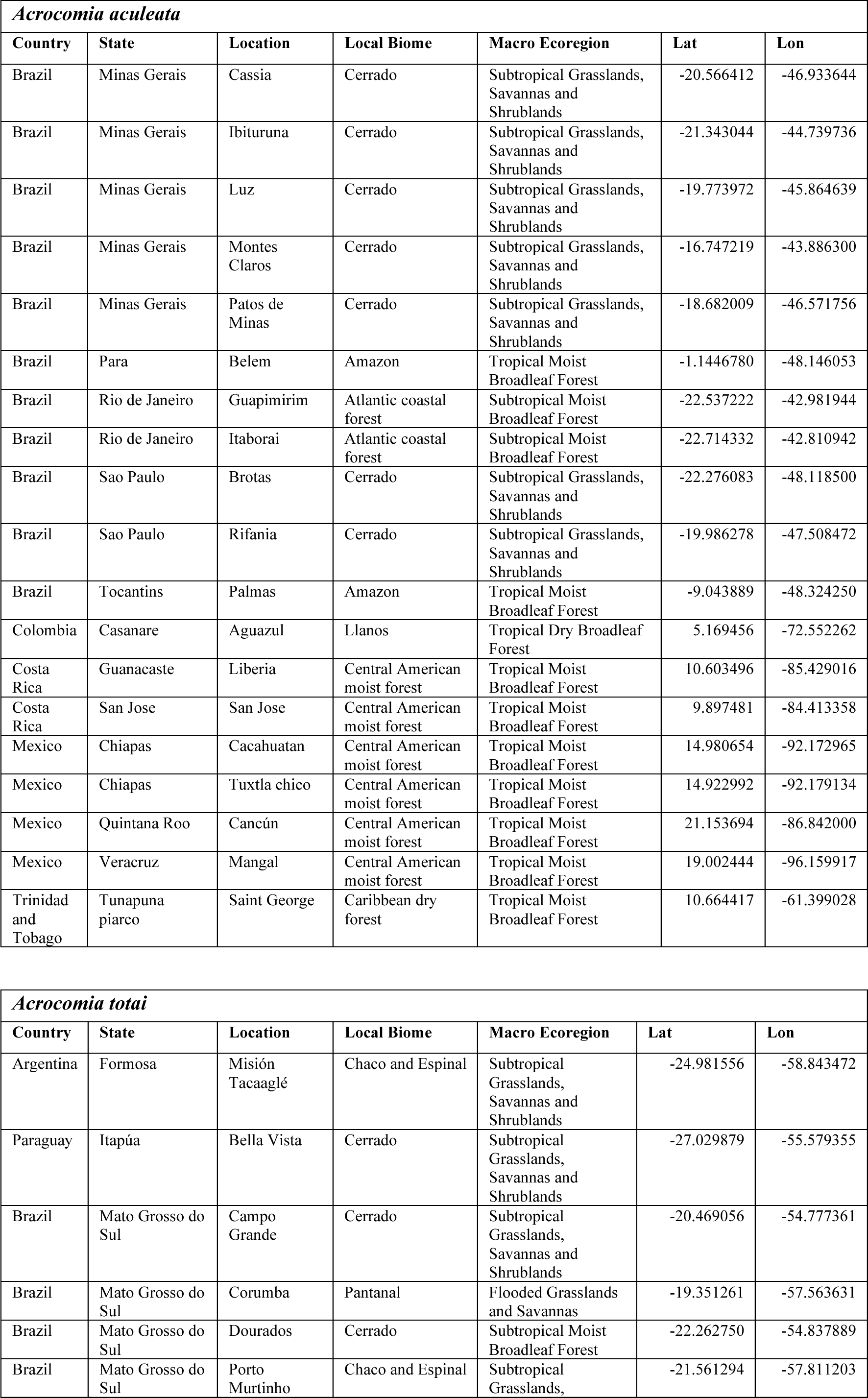
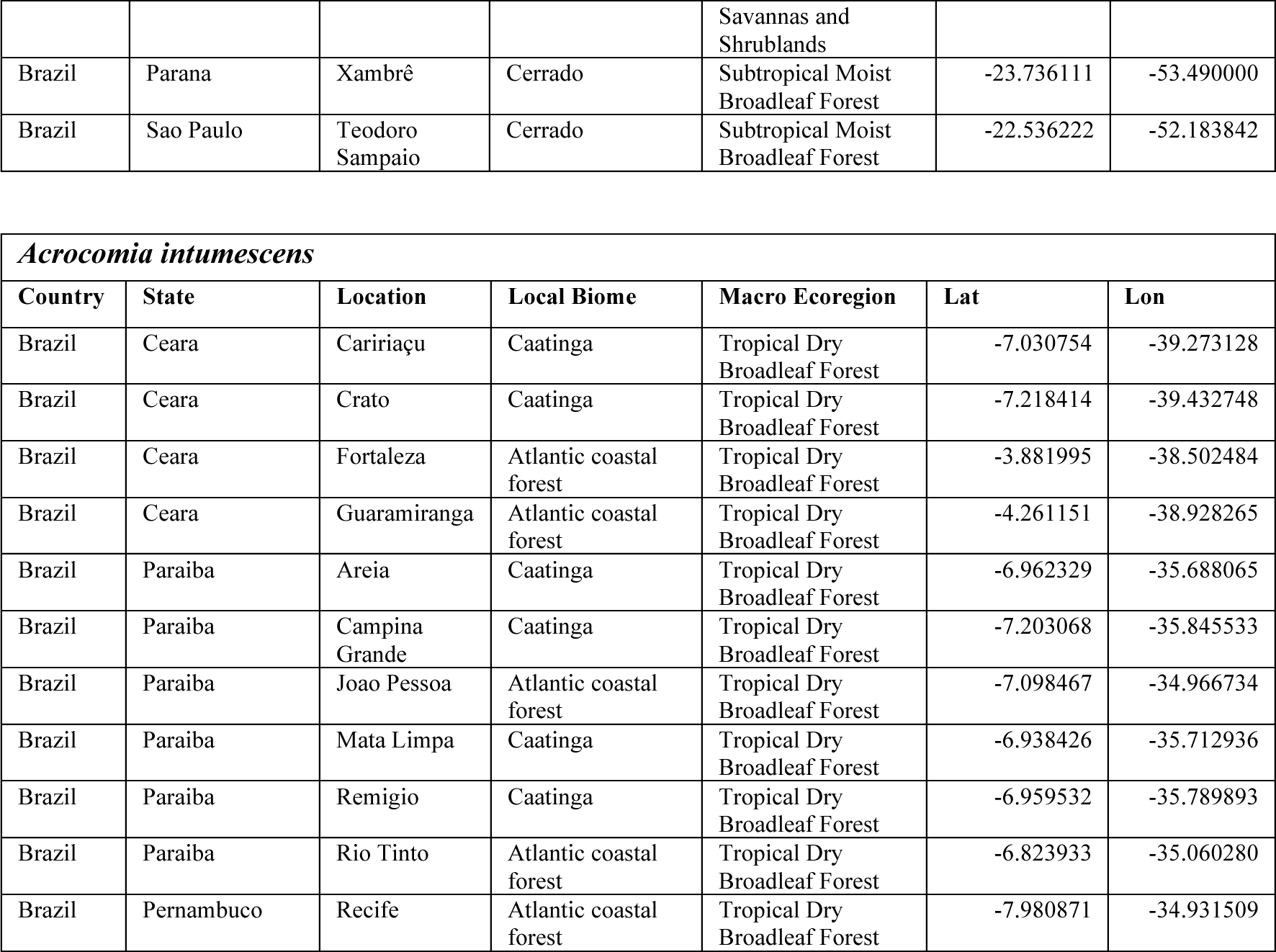
*Acrocomia* species samples. Geographical location and biome of the *Acrocomia* species samples.

### GBS libraries and SNP discovery

Genomic libraries were prepared following the protocol of genotyping-by-sequencing using two restriction enzymes (ddGBS) as described by Poland *et al*. [108]. The combination of *NsiI* and *MspI* (New England Biolabs) was used for the libraries of *A. aculeata* and *A. totai*, while the combination of *NsiI* and *MseI* (New England Biolabs) was used for *A. intumescens*. Two libraries were prepared mixing *A. aculeata* and *A. totai* samples, each using a 96-plex set of *NsiI* barcode adapters and a common *MspI* adapter. Another two libraries were prepared for *A. intumescens*, each using a 96-plex set of *NsiI* barcode adapters and a common *MseI* adapter. The ddGBS libraries were quantified through RT-PCR on the CFX 384 Touch Real Time PCR (BioRad) equipment using a KAPA Library Quantification kit (KAPA Biosystems, cat. KK4824), and the fragments’ profiles were inspected using the Agilent DNA 1000 Kit on a 2100 Bioanalyzer (Agilent Technologies). The libraries of *A. aculeata* and *A. totai* were sequenced on two separate runs in an Illumina NextSeq500, with single-end and 150 bp configurations. The two libraries of *A. intumescens* were sequenced on a single run, but in separate lanes, in an Illumina HiSeq3000, with single-end and 101 bp configurations.

The general sequencing quality and the presence of adapters were assessed with FASTQC [109]. The SNP discovery was performed separately for each species following the *de novo* pipeline of the program Stacks v.1.42 [110] with similar filtering criteria. Due to the presence of adapters, sequences were trimmed to 90 bp for *A. aculeata* and *A. totai*, and 80 pb *A. intumescens*. Trimming, quality control (removal of sequences with uncalled bases and with Phred scores <10), and demultiplexing were performed with the module *process_radtags*. For each sample, loci were assembled using the module *ustacks* considering minimum sequencing depth (-m) of 3, and distance between reads from the same loci (-N) of 2. A catalog of loci across samples was built with the module *cstacks*, considering the distance between locus (-n) of 2, and loci of the samples were compared to the catalog using *sstacks*. The loci with lower probabilities (-lnl_lim 10) were discarded using the module *rxstacks*. For the three species, candidate SNPs were identified using the module *populations* considering a minimum depth of 3, minor allele frequency of 0.01, the presence of SNP in at least 75% of the samples in each of 14 (*A. aculeata*), 8 (*A. totai*), or 10 (*A. intumescens*) sampling locations. To avoid explicit linkage only a single SNP was retained per GBS locus. Quality metrics (mean sequencing depth per locus and per sample, percentage of missing data per sample) were assessed with VCFTools [111]. Additionally, samples with more than 50% of missing genotypes were removed from the final data set, resulting in 69 samples for *A.aculeata*, 40 for *A. totai*, and 131 for *A. intumescens*.

### Detection of putative signatures of selection

Putative signatures of selection were investigated using complimentary approaches [66,112] either based on significant deviations of *F_ST_* estimates among populations, based on principal component analyses (PCA), or based on environmental association analyses, which identify association between environmental variables and individual markers.

A classic approach to identify outlier SNPs is the identification of loci with extremely high or low values of *F_ST_* (or related statistics) estimated among populations. This method was performed using the FstHet [113], package for R [114] which implements a model similar to FDist2 [115] to construct a neutral envelope for the distribution of *F_ST_*s estimated based on the given data set. The neutral envelope was constructed based on 1,000 bootstraps of the *F_ST_*-analogous betahat statistic [116], which considers variations in sampling sizes across populations. In this analysis, the markers below or above the 95% envelope of betahat estimates were considered as outlier SNPs.

Pcadapt [117] was used to identify SNP markers significantly associated with the genetic structure of the data based on a PCA, without assuming any genetic model. This analysis was performed in the pcadapt package for R [114], retaining the first 3, 4, and 1 K principal components for *A. aculeata*, *A. totai* and *A. intumescens*, respectively (Fig. S1). In this analysis, the SNPs with q-values ≤ 0.1 (corrected p-values for their association with the first K principal components) were considered as outliers.

The Latent Factor Mixed Models (LFMM) analysis [118] was used to assess the correlations of SNP markers with environmental variables obtained in the *WorldClim2* data base [119]. The information recovered from *WorldClim2* refer to 19 bioclimatic variables, of which 11 are primarily related to annual trends of temperature (BIO1 to BIO11) and eight are related to annual trends of precipitation (BIO12 to BIO19). For each species, the variables were extracted for the sampling points, and a Pearson’s correlation test was employed to minimize interdependence among them, and only the variables with a correlation coefficient < 0.8 were retained (Fig. S2). A sparse non-negative matrix factorization (sNMF) analysis [120] was performed to estimate the most likely number of genetic clusters for each species to model the covariation of the subjacent genetic structure in LFMM. Both, sNMF and LFMM were performed with the package LEA [121] for R [114]. For each species, 10 independent repetitions of sNMF simulating from 1 to 10 *K* ancestral groups were performed with 200,000 iterations. The most probable number of *K* = 2 for *A. aculeata* and *A. intumescens*, and *K* = 4 for *A. totai* were estimated according to the cross-entropy estimates of the algorithm (Fig. S3). These numbers were used in subsequent LFMM analyses, which were performed based on 10 repetitions of 50,000 burn-in followed by 100,000 iterations of the algorithm for each species. The generated p-values were corrected considering a false discovery rate (FDR) of 0.1 as threshold for the identification of SNPs significantly associated with the environmental variables.

SNPs were declared as putatively under selection if identified as outliers (or significantly associated to environmental variables) by at least two of the three methods described above. Blast2GO [122] was used to assess similarities between the GBS tags with outlier SNPs and proteins with described putative functional annotations. The similarity between GBS tags in which outlier SNPs were identified and proteins deposited in GenBank was evaluated using blastx with default configurations but restricting the search to Viridiplantae data. Sequences with blastx hits were then screened against the Pfam database to search for protein domains and their associated Gene Ontology (GO) annotations. The putative annotations were summarized based on the GO terms and visually represented with bar plots using the on-line tool WEGO (http://wego.genomics.cn/).

## Authors contributions

JM-M, MIZ, and CAC conceived and coordinated the study. JM-M, BD-H, SAV and CBA led the biodiversity prospecting and conducted the molecular biology assays and Next Generation Sequencing library construction. AA-P performed the SNP calling and conducted genetic statistics. JM-M drafted the manuscript. BD-H, SAV, CAC, AA-P, MIZ, and JBP contributed to the editing of the manuscript. MIZ and CAC contributed to the funding of the research. All authors read and approved the final manuscript.

## Acknowledgements

We acknowledge the financial support from the São Paulo Research Foundation (FAPESP— grants: 2021/10319-0, 2019/20307-0, 2014/23591-7) and the National Council for Scientific and Technological Development (CNPq—grant #431226/2018-0). We also thank the Coordination for the Improvement of Higher Education Personnel of Brazil (CAPES) for the granted scholarship Ph.D. to JM-M. Also thanks the São Paulo Research Foundation for a post-doctoral scholarship to AA-P (FAPESP 2018/00036-9).

## Supporting information

**Fig. S1 Principal component analysis (PCA) performed for outlier detection using pcadapt for *Acrocomia aculeata*, *A. totai* and *A. intumescens*.** Top row: scree plots of the explained variance (y-axis) retained in each principal component (PC) (x-axis). The number of retained PCs was chosen as the first inflection point (Cattell’s rule), where the amount of genetic variation added by successive PCs reaches a plateau. Bottom row: the associated scatter plots of the first two PCs.

**Fig. S2 bioclimatic variables recovered from WorldClim2 and extracted for the sampling points of *Acrocomia aculeata*, *A. totai* and *A. intumescens*.** Groups of intercorrelated variables highlighted within red boxes (Pearson’s correlation coefficients > 0.8).

**Fig. S3 Sparse non-negative matrix factorization (sNMF) performed for outlier detection using Latent Factor Mixed Models (LFMM) for *Acrocomia aculeata*, *A. totai* and *A. intumescens*.** Left: plot of the cross-entropy estimates for each number of simulated ancestral populations. Right: Bar plots representing the sNMF ancestry coefficients across samples from different biogeographic group in each species. Each sample is represented by a bar and different shade colors represent their associated ancestry proportion from distinct genetic groups. Acronyms follow Table 2: CMF Central American Moist Forest; LLA Llanos; CDF Caribbean Dry Forest; ACF Atlantic Coastal Forest; CAA Caatinga; AMA Amazon; CER Cerrado; PAN Pantanal; CHE Chaco and Espinal.

