## Supplementary figures and images for "Genetic variations associated with adaptation processes in Acrocomia palms: A comparative study across the Neotropic for future crop improvement"

### Fig. S1

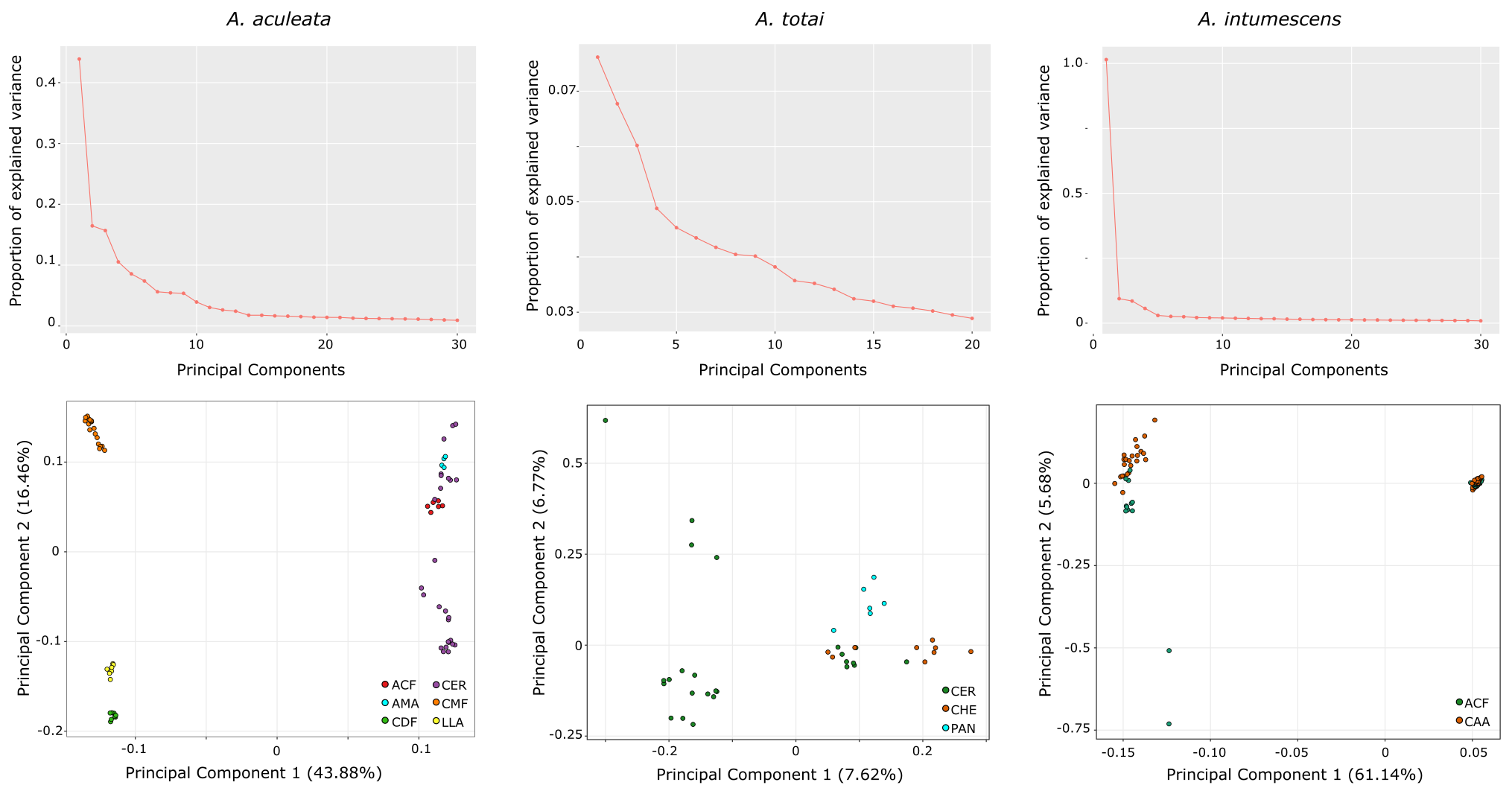

### Fig. S2

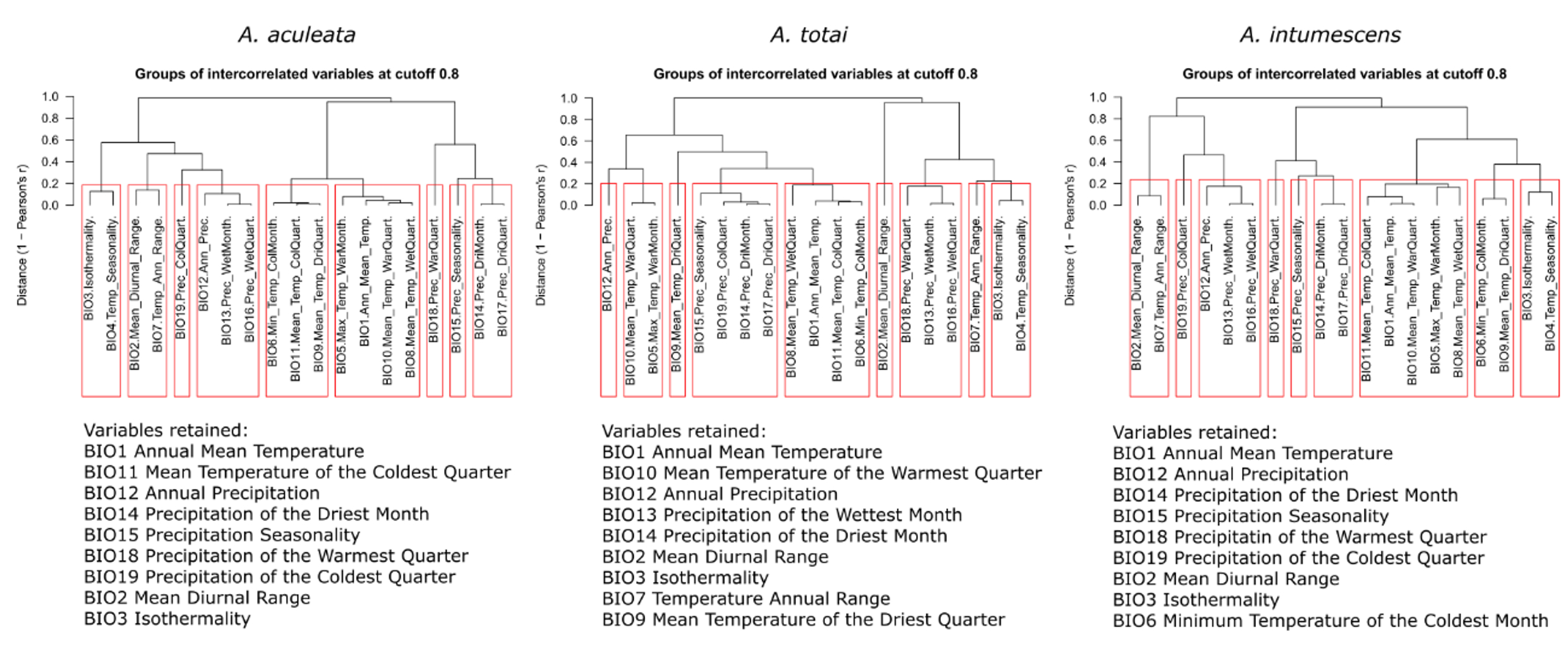

### Fig. S3

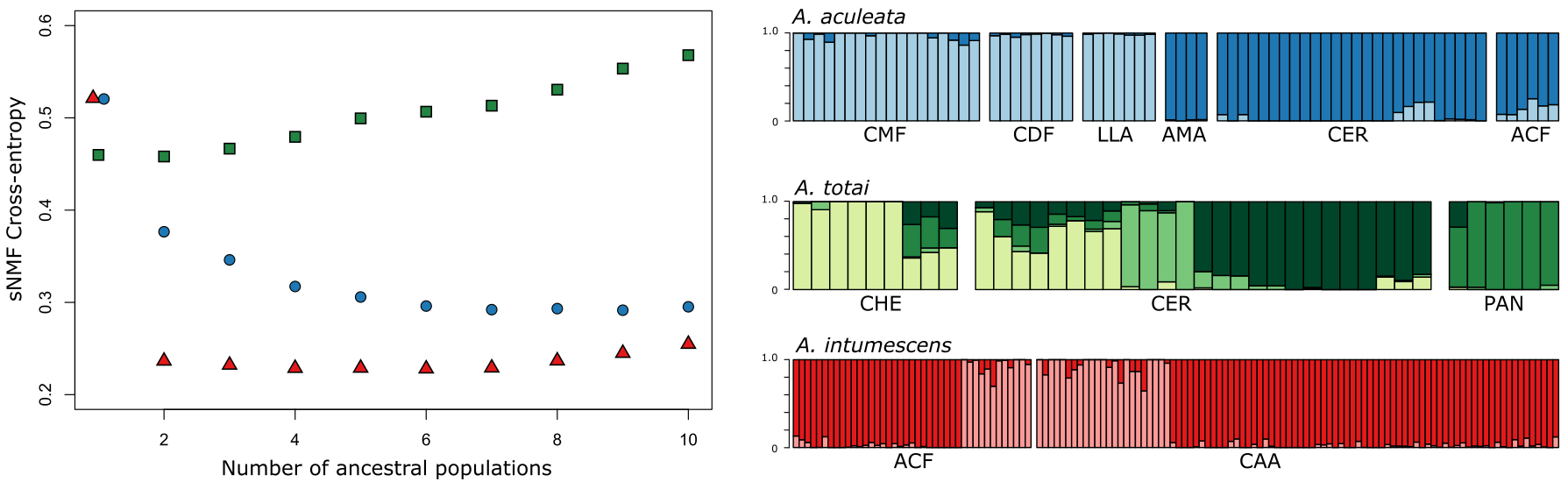
